# One-cell Doubling Evaluation by Living Arrays of Yeast, ODELAY!

**DOI:** 10.1101/069724

**Authors:** Thurston Herricks, David J. Dilworth, Fred D. Mast, Song Li, Jennifer J. Smith, Alexander V. Ratushny, John D. Aitchison

## Abstract

Cell growth is a complex phenotype widely used in systems biology to gauge the impact of genetic and environmental perturbations. Due to the magnitude of genome-wide studies, resolution is often sacrificed in favor of throughput, creating a demand for scalable, time-resolved, quantitative methods of growth assessment. We present ODELAY (One-cell Doubling Evaluation by Living Arrays of Yeast), an automated and scalable growth analysis platform. High measurement density and single cell resolution provide a powerful tool for large-scale multiparameter growth analysis based on the modeling of microcolony expansion on solid media. Pioneered in yeast but applicable to other colony forming organisms, ODELAY extracts the three key growth parameters (lag time, doubling time, and carrying capacity) that define microcolony expansion from single cells, simultaneously permitting the assessment of population heterogeneity. The utility of ODELAY is illustrated using yeast mutants, revealing a spectrum of phenotypes arising from single and combinatorial growth parameter perturbations.

## INTRODUCTION

Growth is a well-established, sensitive metric of cellular fitness that is widely used to interrogate genetic and environmental interactions. The most basic models of microorganism population expansion over time consist of three distinct phases – lag phase, log phase and stationary phase (Monod, 1949). Each phase is defined by a specific parameter that uniquely contributes to overall fitness. Lag phase, defined by lag time, is the period after initial inoculation wherein little to no growth is observed. Following acclimation, the population enters log phase and expands exponentially at a constant, maximal rate defined by the doubling time. Finally, a rapid cessation of growth is observed as the population enters stationary phase, having reached its maximum attainable level defined by the carrying capacity. By virtue of its linear nature during exponential growth, the log plot of population number versus time has classically been employed to extract the three key growth parameters. Lag time is the period up to the attainment of linearity of the log-plot, doubling time is inversely proportional to the slope of the linear region of the log-plot and carrying capacity is the maximum population size when the slope of the log plot approaches zero.

In light of its relatively well-understood cell biology and genetic tractability, baker’s yeast, *Saccharomyces cerevisiae*, is a model organism commonly exploited to elucidate genetic and environmental interactions on a genome-wide scale. Many methods of assessing yeast strain growth characteristics have been described and most employ liquid culturing (Breslow et al., 2008; Bryan et al., 2010; Godin et al., 2010; Kortmann et al., 2009; Murakami and Kaeberlein, 2009; Sun et al., 2010; Tucker and Fields, 2004; Winzeler et al., 1999). These include direct measurements, such as cell counting and flow cytometry, and indirect measurements, the most common being the turbidity of the growth media measured by absorbance of 600 nm light (OD_600_). Dynamic range limitations associated with many of these methods render them unable to assess all three growth parameters within a single experimental run; thus, analyses are often restricted to only one growth parameter, most commonly doubling time. Furthermore, difficulties associated with maintaining low volume yeast cultures in suspension at high densities limit the throughput of many liquid growth analysis techniques (Kortmann et al., 2009).

The shortcomings inherent to yeast liquid culture analyses have made it commonplace to employ cell spotting as a proxy for strain growth. Cell spotting assays range from biofilm analysis, in which a population of cells is delivered as a patch onto the surface of solid media, to serial dilution analysis, wherein single colonies are obtained (Lawless et al., 2010; Memarian et al., 2007; Shah et al., 2007). While these methods are universally accepted, there are major caveats to their use. Foremost, despite the demonstration of dynamical growth assessment for populations of cells through the analysis of biofilm intensity on solid media (Shah et al., 2007), most large-scale fitness analyses are assessed from a single time point (Baryshnikova et al., 2010; Collins et al., 2006; Costanzo et al., 2010). The lack of temporal resolution makes it impossible to deconvolve the different stages of population growth and, therefore, apparent differences in fitness cannot be unequivocally attributed to the classically defined growth parameters of doubling time, lag time and carrying capacity.

More recently, flatbed scanners have used to periodically image growing biofilms (Levin-Reisman et al., 2010). However, most flat-bed scanners have an optical resolution slightly greater than 5 μm per pixel, which cannot reliably image individual cells. The lack of resolution limits the initial measurement to a relatively late stage in population development, when colonies/biofilms can be clearly resolved. Consequently, lag time is not directly observed and edge effects and other local competition artifacts are present. In the case of the widely used synthetic genetic array (SGA), epistatic miniarray profile (E-MAP) and SCANlag methods, effects of some confounding factors are corrected by the latest generation of analytical tools; however, given that multiple data sets involving many query strains are required to normalize for batch effects (Baryshnikova et al., 2010), sensitivity is proportional to the scale of the study using these methods, which limits their tactical utility.

In this work, we present a platform capable of high-density measurements of lag times prior to the attainment of doubling times during exponential growth, and carrying capacities at stationary phase through time course microscopy-based imaging of microcolonies growing on solid media. Because each microcolony is seeded from one to a few cells and hundreds of microcolonies can be analyzed for each strain, population heterogeneity of the three growth parameters can be assessed on a strain by strain basis. Through increased sensitivity and the potential for growth parameter profiling, the enhanced resolution afforded by this novel method of multiparameter fitness assessment can facilitate the generation and/or refinement of gene-gene and gene-environment interaction networks for yeast and other colony forming organisms.

## MATERIALS AND METHODS

### Yeast strains and growth conditions

Unless otherwise specified, all experiments were performed at 30°C temperature using rich growth media, YEPD, [1 % w/v yeast extract (BD), 2 % w/v peptone (BD), 2 % w/v dextrose (BD)]. Galactose growth media contained 2 % w/v galactose (Acros) in place of glucose and solid media contained 2 % w/v agar (BD) for cell spotting assays or 1.0 % w/v agarose (Invitrogen) for ODELAY analyses. *S. cerevisiae* strains used in this study are listed in Supplementary Table 1 online. All strains have been previously described^9,20^.

**Table 1:**
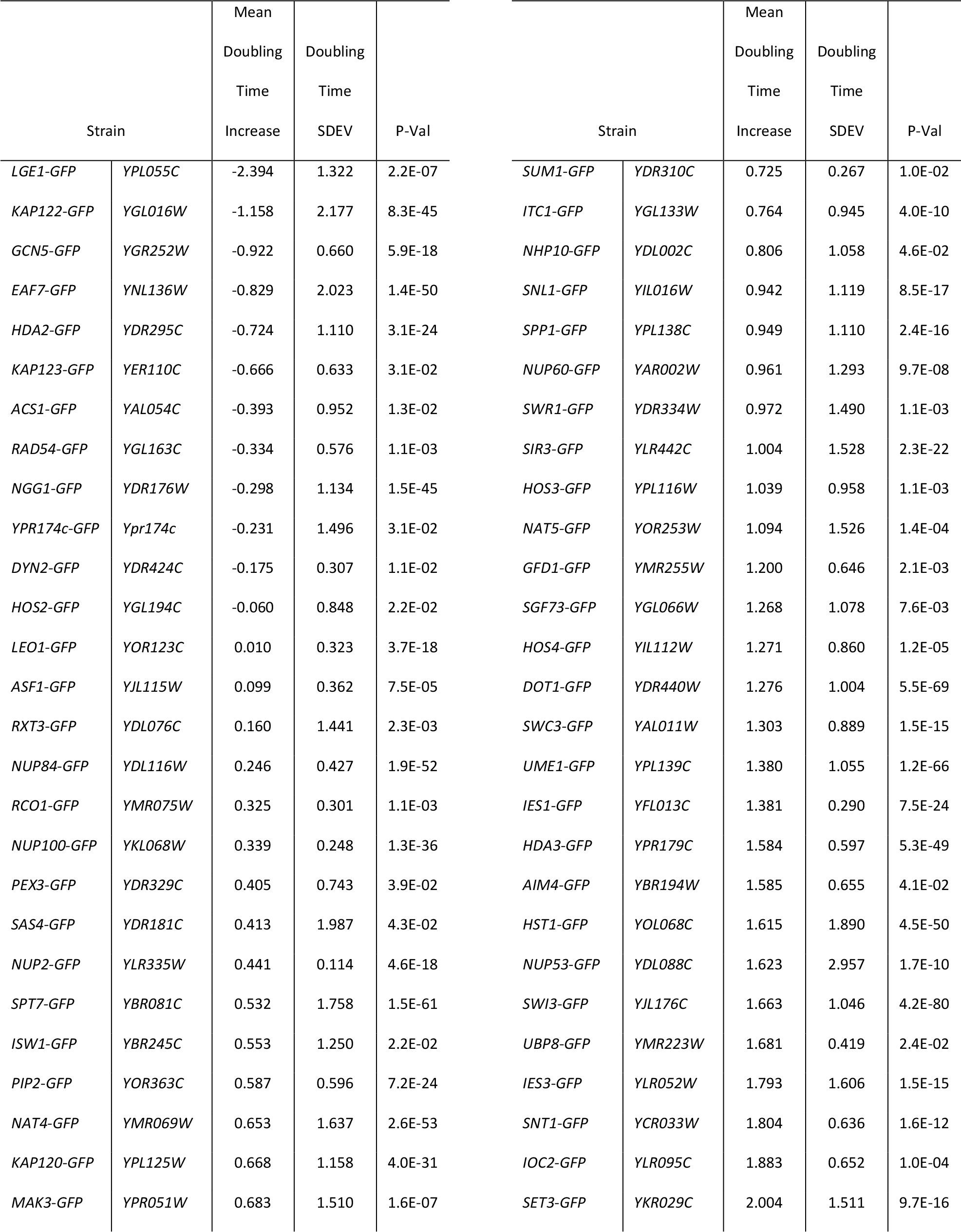
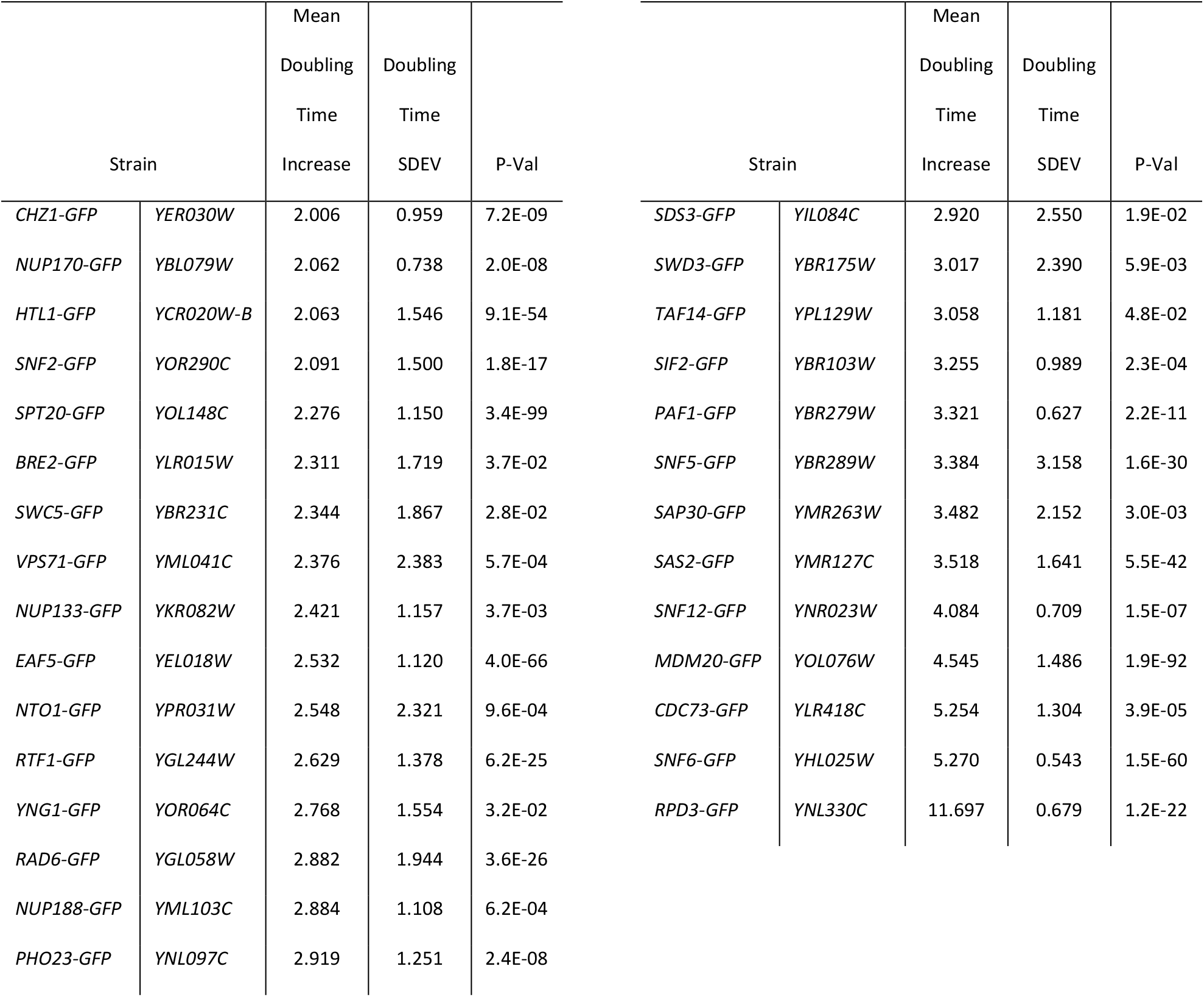
GFP-tagged library mutants with significant growth difference over wild type.

### ODELAY Culture Preparation

Stringent culture conditions were required for reproducible growth phenotypes. 220 μL yeast cultures were inoculated in 96 well flat bottom plates (Corning Costar) and grown overnight. Cultures were then diluted 1:11 and optical densities read using a Synergy H4 plate reader. Individual wells were then diluted to a density of 0.09 OD and the culture grown for 6 hours to ensure all strains were in exponential phase. The cultures were again measured using the plate reader and then diluted to 0.01 OD. The 96 well plate containing the prepared cultures was then sonicated for 30 second in an ice-cold water ultrasonic bath to dissociate cell clusters.

### ODELAY Slide preparation and yeast array setup

Growth media was prepared as a 1:1:8 mixture of the following sterile stock solutions, respectively, 10X YEP (10 % w/v yeast extract and 20 % w/v peptone), 20 % w/v carbon source (glucose or galactose) and 1.33 % w/v agarose in water. Typically, a 150 mL volume of 1.33 % agarose stock was prepared, divided into 15 mL aliquots in 50 mL conical bottom tubes and stored at 4 °C until use. Agarose aliquots with 2 mL 10X YEP and 2 mL 20% carbon source were placed in rapidly boiling water for 20 minutes to completely melt the agarose gel. Water lost to evaporation was replaced by weighing the conical tube before and after boiling, yielding a final growth substrate containing 1 % yeast extract, 2 % peptone, 2 % carbon source and 1 % agarose. The molten solution was poured into custom molds that formed 2 mm slabs of agar supported by 50 mm by 75 mm by 0.1-inch glass slides (Fisher Scientific). The apparatus was allowed to cool to room temperature and, after careful separation of the glass slides, the agar plates were equilibrated overnight in a humidified chamber. Careful separation of the glass slides was critical as any mechanical deformation of the agar altered the lag time and doubling time of cultures in the regions deformated. The following day, yeast in exponential liquid culture, diluted to an OD_600_ of ∼0.01, were spotted onto agarose slabs using a Matrix Hydra DT fluidics robot (ThermoScientific). Slides were air dried for ∼3-5 minutes and then placed inside a microscope equipped with a humidified environmental chamber maintained at 30°C.

### ODELAY Image acquisition time course

Bright field images were captured using a Leica DMI6000 microscope (Leica) equipped with a 10X objective. Images were recorded by a Hammamasu ORCA Flash 4.0. The microscope stage movements and camera were controlled by a custom MATLAB graphical user interface using the Micromanager Core API version 1.4(Edelstein et al., 2014). MATLAB scripts controlled the stage to predefined positions. A custom autofocus routine found focus at the center of each spot by maximizing the image’s focus score using utilizing the Laplacian variance function(Pertuz et al., 2013). After focus was found a 3 × 3 tiled image was recorded which covered a 9 mm^2^ area of the agar. These steps were repeated on each of the 96 spotted strains in either 30 minute or 1 hour increments for 48 hours.

### Automated ODELAY image analysis

Image acquisition and panorama stitching were performed using MATLAB scripts. Briefly images were stitched using a method based on FFT phases(Preibisch et al., 2009). A threshold of the stitched images was calculated by taking histograms of a subdivided image and finding the maximum intensities of 100 regions within the subdivided images. This threshold was used to binarize images and colony area was data quantified using MATLAB functions. The log_2_ of colony area was plotted versus time and colony area fit to a parameterized version of the Gompertz function (*f*)(Gompertz, 1825; Zwietering et al., 1990),

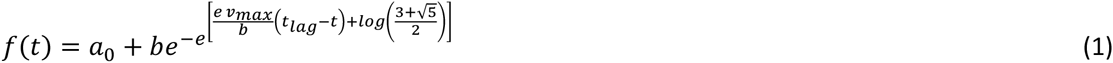

Where *a*_*o*_ and *b* are parameters that represent the initial size and final saturation of the colonies; *v*_*max*_ maximum growth velocity, and *t_lag_* colony lag time. Growth parameters were solved for directly. The *gompertzFit* routine calculates an initial estimate of the Gompertz function (1) using a coarse grid optimization and then attempts to find a constrained minimum of the function (1) at this initial estimate using the *fmincon* MATLAB function. In order to proceed to curve fitting, colonies must be matched at 5 or more time points through the monitored time course. In addition, colonies that do not exhibit at least a doubling in area are eliminated from curve fitting. This is achieved by only fitting data for which the maximum observed cross-sectional area of each tracked object is at least two-fold greater than the object’s measured cross-sectional area at the first time-point. Doubling time (*t_d_*) is calculated as follows:

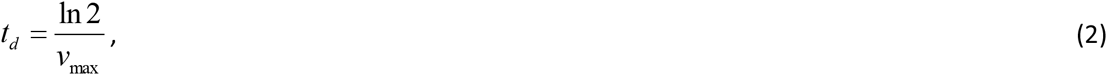

Where *v*_*max*_ is the point at which the growth rate, *f’(t)*, reaches maximum (achieved at *f’’(t)*= 0). Lag time (*t_lag_*) is defined as the time to reach maximum growth acceleration, *a_max_*, where *f’’(t)*is greatest (achieved at the lower value of the two solutions to *f’’’(t)*= 0). The carrying capacity (*K*), in pixel area, represents the cross-sectional area of the base of the modeled microcolony projected to stationary phase ( *f(t)* as *t*→∞) and is calculated as follows:

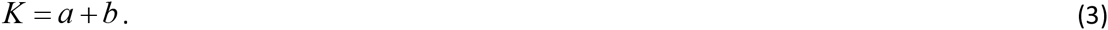

### BioScreen doubling time determination

Automated optical density measurements of yeast cultures were obtained using a BioScreen C (Growth Curves USA) using manufacturer’s suggested protocols with the exception that culture volume was reduced to 200 μL to prevent artifacts arising from liquid splashing onto the plate lid during maximal agitation. A starting OD_600_ of 0.05 was utilized in order to ensure that cultures were in exponential phase once they entered the empirically determined linear range of the instrument. Growth curves were fit using the *gompertzFitBioScreen* function with is identical to the *gompertzFit* function except optimized for the range of OD_600_ values obtained from the Bioscreen C instead of observed area.

## RESULTS AND DISCUSSION

### Development of an automated scalable solid-phase doubling time estimation platform

An ideal solid-phase, time-resolved, growth analysis platform would allow for high sample density and be amenable to automated data acquisition and processing. The optimized method, which we have termed ODELAY for <u>O</u>ne-cell <u>D</u>oubling <u>E</u>valuation by <u>L</u>iving <u>A</u>rrays of <u>Y</u>east, is depicted schematically (Fig. 1) and consists of four stages: spotting of ordered arrays of live yeast onto thin beds of growth substrate on a glass slide support (Fig. 1A); periodic bright field image acquisition over a user-specified time course (Fig 1B); processing of raw bright field data to extract microcolony cross-sectional area data (Fig. 1C) and post-processing calculation of growth parameters for each individual microcolony within each spot (Fig. 1D). ODELAY is applicable to a wide range of growth substrates and incubation temperatures and is highly scalable, as it can analyze 10^5^ to 10^6^ individual microcolonies per experiment.

**Figure 1.**
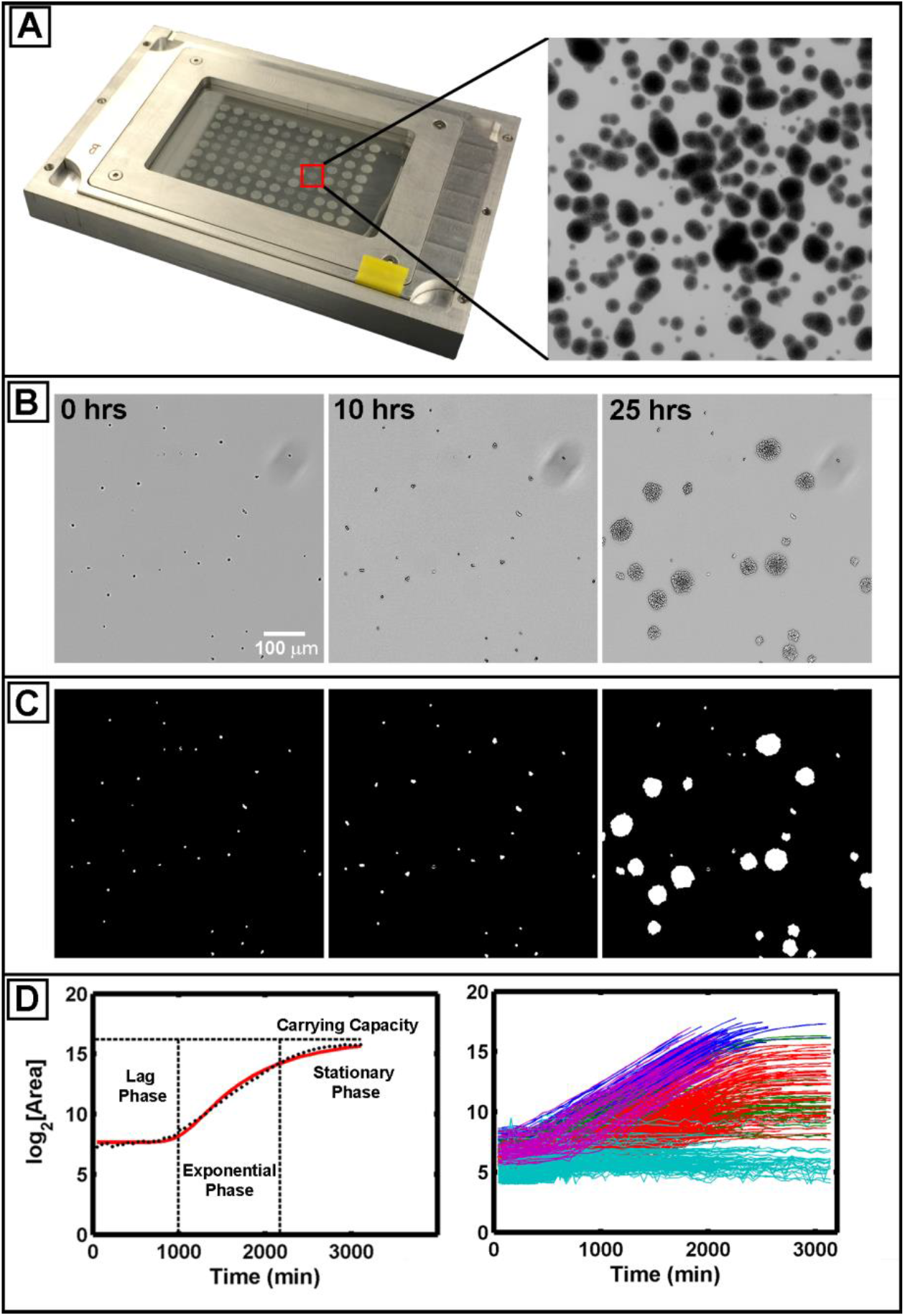
One-cell Doubling Evaluation by Living Arrays of Yeast (ODELAY). Solid-phase growth parameters are extracted by collecting time course image of growing colonies (1A and 1B). Colony areas are measured from thresholded and binarized images from the time course image series (1C). Colonies seeded from single yeast cells are tracked over time and the log2(Area) is used to fit a parameterized version of the Gompertz function (1D left). A wild-type yeast strain (BY4741) was pre-grown to saturation with glucose as a carbon source and then assayed on galactose-containing agar. The resulting to represent each cluster (1D right).

ODELAY consists of an automated pipeline that encompasses acquisition and processing of images, identification and measurement of microcolonies at each time point, matching of microcolonies through time and extrapolation of growth parameters from growth curves. This current platform employs theoretical approximation of ODELAY growth curves using the Gompertz function as an unsupervised method to extract growth parameters(Gompertz, 1825; Preibisch et al., 2009). All files required for execution of automated ODELAY analysis, as well as a demonstrative data set, are available as Supplementary Files online.

### Determination of growth parameters by ODELAY

First, data are acquired, and then growth parameters of doubling time, lag time and carrying capacity are determined by directly fitting a parameterized version of the Gompertz function (Eq. 1). For data acquisition, the first time point would ideally be acquired immediately after spotting onto agar at the desired growth temperature, but for practical purposes, the starting time is when the cells are spotted at room temperature on the solid substrate. The plate is then transferred to an environmentally controlled chamber and growing colonies are tracked until they merge with their neighbors. The time required for colonies to merge is therefore related to the initial cell density and the ultimate carrying capacity of adjacent colonies. While many colonies merge before carrying capacities are observed, ODELAY will still estimate carrying capacity as long as a sufficient number of data points are collected after maximum growth velocity is achieved. This is a feature of the Gompertz function’s symmetry about maximum growth velocity, which permits fair estimation of carrying capacity even when it is not directly measured. Note that caution should be exercised when examining phenotypes associated with increased carrying capacity because the Gompertz function may not accurately estimate all possible outcomes.

### Comparison of ODELAY to established methods

We directly compared doubling times and lag times calculated by multiple ODELAY population measurements to liquid culture OD_600_ measurements made using the BioScreen C instrument for both fast and slow growing strains taken from the MATα yeast deletion library (Winzeler et al., 1999) (Fig. 2A). Population doubling times and precision of this measurement across replicates were roughly comparable between the two platforms (Fig. 2A). Measured doubling times for 140 yeast strains correlated well between the two platforms with a Pearson coefficient of 0.76 and Spearman coefficient of 0.70 (Fig. 2B). Lag times showed less agreement, likely due to the liquid versus solid culture medium, and the lack of sensitivity of optical density measurements at low cell concentrations. In addition, unlike BioScreen, ODELAY identified slow growing outliers because microcolony growth curves are derived from single cells. In contrast, liquid culture OD_600_ curves measure an aggregate of all cells in a population, and therefore are not sensitive to the contribution of individual cell’s.

**Figure 2.**
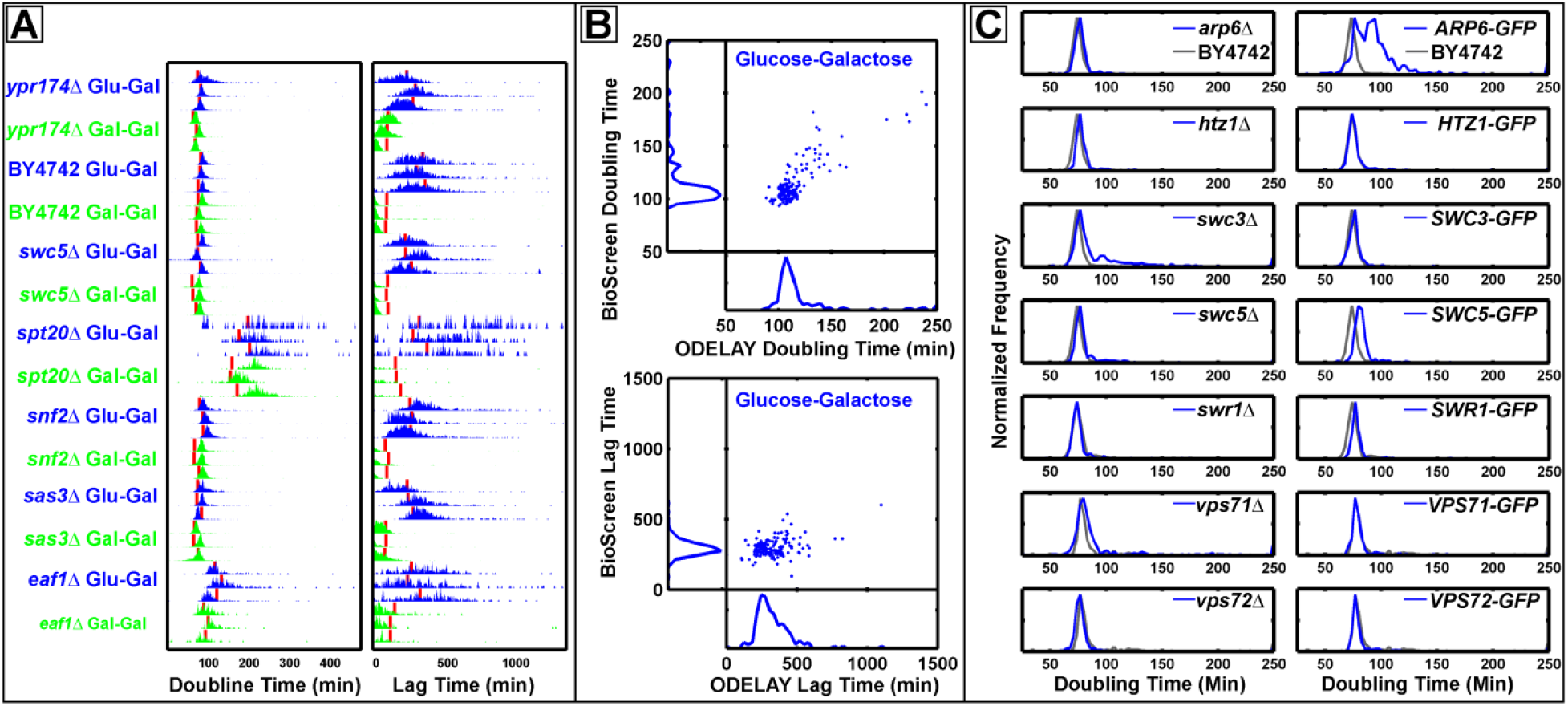
Complex phenotypes observed by ODELAY. Comparisons of doubling times and lag times for repeated measurements (2A). Red lines indicate Bioscreen C results. Median ODELAY measurements show good agreement with BioScreen C measurements in doubling time but less so in lag time (2B) Population histograms of doubling time from SWR1 complex deletion and GFP-tagged strains (blue) with comparison to the parent strain BY4742 (grey) (2C). Heterogeneity in doubling times is observed in strains *swc3* and ARP6-GFP while *arp6* and SWC3-GFP appear similar to the parent strain BY4742.

Microcolony convergence is the limiting factor of ODELAY’s dynamic range, which can be controlled by altering the initial cell density obtained when spotting yeast cultures. In contrast, the dynamic range of liquid culture measurements is limited by either the nutrient capacity of the media or the linear range of the density sensor. Due to differences in strain doubling times, the dynamic range is best defined by the total number of doublings required to reach the upper limit starting from a single cell. At optimal seed density (∼25 – 50 cells/mm^2^), the dynamic range of ODELAY is 8 - 12 doublings – from a single cell up to 250 or as many as 4000 cells, which compares favorably to a dynamic range of 3 – 5 doublings attainable by most currently available technologies.

In traditional biofilm assays, the size of a colony is dependent on the number of viable individuals contributing to the colony population, the number of doublings these cells have undergone, the amount of nutrients present, and the ability of the colony to transport nutrients to its reproducing members. The contribution of individuals to the overall colony size is not distinguished by traditional methods such as liquid based assays or spot based assays. In contrast, ODELAY tracks individual cells forming into colonies and can quantify population heterogeneity that other methods cannot resolve.

### Example applications of ODELAY

#### Identification of doubling time phenotypes

To illustrate the ability of ODELAY to compare population heterogeneity of growth phenotypes between strains such that features of the population distributions may be evaluated we focused on members of the SWR1 complex of chromatin modifiers (Fig. 2C). Chromatin modification is one way for the emergence of epigenetic differences that can manifest as heterogeneity within isogenic populations. We observed population heterogeneity in two strains *swc3Δ* and *ARP6-GFP* (Fig. 2C) but not in their respective GFP tagged or deletion mutant. Population heterogeneity has been observed before in SWR1 deletion strains when measuring *POT1-GFP* expression during a carbon source switch from glucose to oleic acid(Knijnenburg et al., 2011). In that instance, deletion of other members of the SWR1 complex induced bimodal expression of *POT1-GFP*. Here, the bimodality of growth phenotypes emerged from cells grown strictly on glucose media and without any stimulation from a change in carbon source. This observation demonstrates that ODELAY readily detects subpopulations of cells present in standard culture of deletion and GFP fusion strains.

#### Identification of lag time phenotype

Through ODELAY analysis, outliers with highly variable, expanded or contracted lag periods can be identified by assessing the distribution of lag times for microcolonies of a given strain (lag time variability) as well as relative lag between tested strains. To demonstrate the quantification of lag time by ODELAY, we exploited the well-studied and highly regulated response of yeast to a carbon source shift from its preferred source, glucose, to an alternative source, galactose(Ideker, 2004; Sellick et al., 2008). This shift is characterized by a lag phase, during which, the normally repressed galactose utilization machinery, including the galactose transporter, is induced. Exponentially growing yeast preconditioned in either glucose or galactose liquid medium were spotted onto solid media containing galactose and analyzed by ODELAY (Fig. 4A). Cells preconditioned in galactose media exhibited a highly synchronized response characterized by short lag times. In contrast, more pronounced and variable lag times were observed for cells that were not primed for growth in galactose. Once glucose grown cells acclimated to the shift to galactose and entered exponential phase, they doubled at rates similar to those observed for galactose preconditioned cells.

As with heterogeneity of doubling times, ODELAY enables the detection of heterogeneity in lag times. To demonstrate the utility of ODELAY in assessing population heterogeneity of lag times, we compared growth parameters of a wild-type yeast strain (BY4742) in galactose-containing medium after pre-growth in glucose media for differing amounts of time (Fig. 3A). We staggered seeding of cultures such that cells were pre-grown in glucose media for 3, 6, 24 and 48 hours (Fig. 3B). The resulting cultures were then spotted on galactose media and their growth phenotypes observed (Fig. 3C). Not only did lag time correlate with the length of time that yeast was cultured in glucose but also colony-to-colony variation in lag times increased for the longer incubation times. This example demonstrates ODELAY’s ability to capture the effects of environmental perturbations on population heterogeneity, a feature which is difficult to distinguish using other solid-media growth assays.

**Figure 3.**
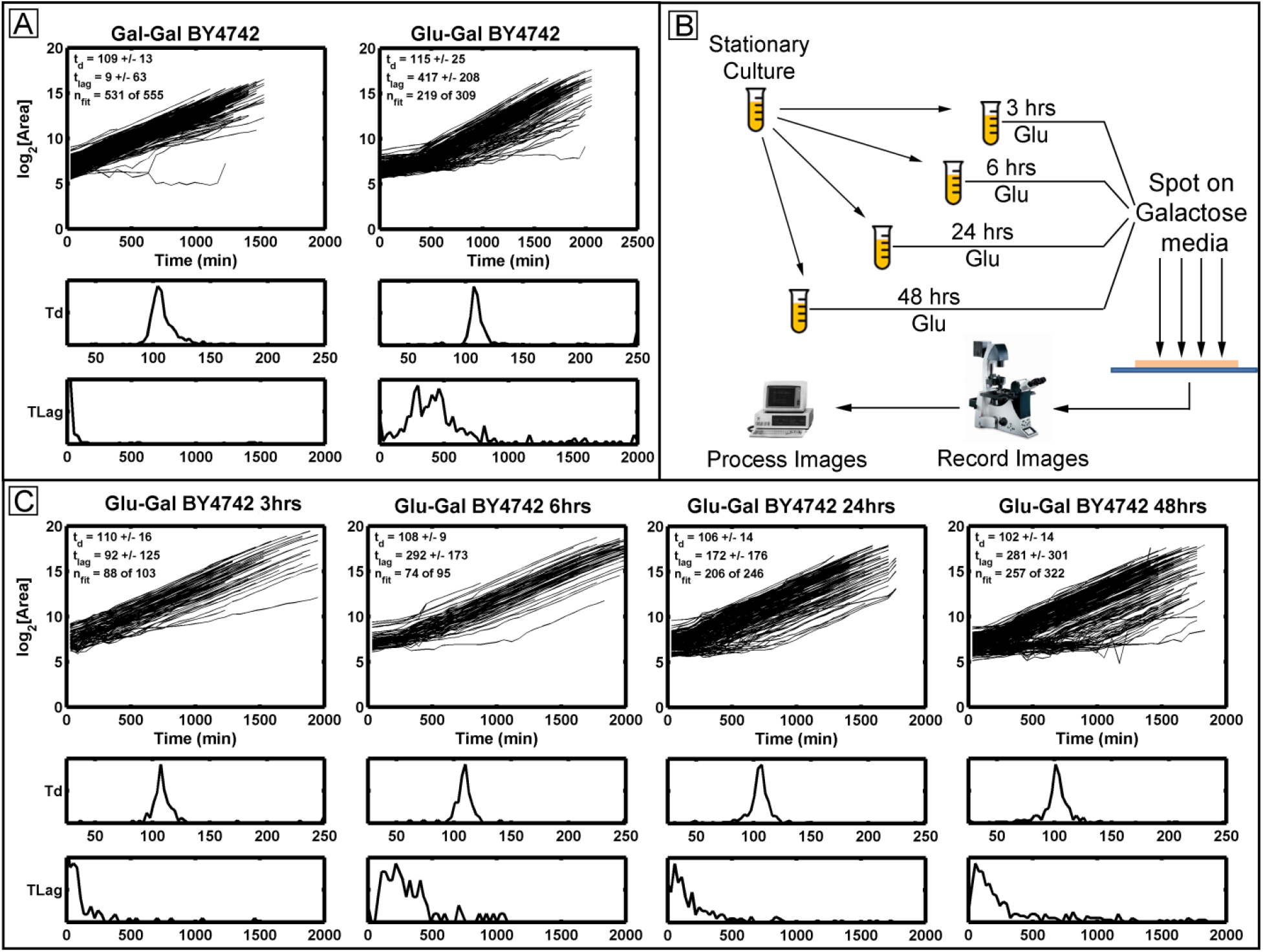
Observing lag time after a carbon source switch: Growth curves and histograms depicting lag time (TLag) and doubling time (Td) for wild type yeast (BY4742) pre-grown in galactose (left) or glucose (right) and then spotted on galactose media (A). Differing growth phenotypes are observed from samples of a wild type yeast strain taken from the same culture at multiple times after seeding a source culture (B and C). Histograms of the doubling time (Td) and lag time (TLag) are depicted below each set of growth curves. Note the changes in the lag time distributions as the source culture is aged. This demonstrates ODELAY’s utility and sensitivity to culture conditions of yeast.

#### Large-scale multiparameter analyses with ODELAY

A strength of the ODELAY platform is to extract doubling times and lag times for populations of cells growing on solid media in a high-throughput manner. To demonstrate multiparameter growth rate analysis by ODELAY, we assayed a collection of 140 strains that contained gene deletions of transcription factors, transcriptional regulators, and nuclear transport factors including nucleoporins and karyopherins. The genes selected were previously associated with regulating the response to a carbon source shift(Winzeler et al., 1999; Knijnenburg et al., 2011; Van de Vosse et al., 2011; Aitchison and Rout, 2000).

For the deletion strains, we quantified colony doubling times, lag times and estimated carrying capacities during a carbon source switch from glucose to galactose, using galactose to galactose transition as a control. This rich multivariate dataset underscores how ODELAY can reveal complex and heterogeneous growth phenotypes of populations of individual cells growing into colonies (Fig. 4). Strains with noticeably strong increases in doubling time include *dot1Δ*, *htl1Δ*, *eaf5Δ*, *eaf7Δ*, and *spt20Δ*. Of these five examples, only *spt20Δ* had been reported to have reduced growth rate on galactose media(Roberts and Winston, 1996).

**Figure 4.**
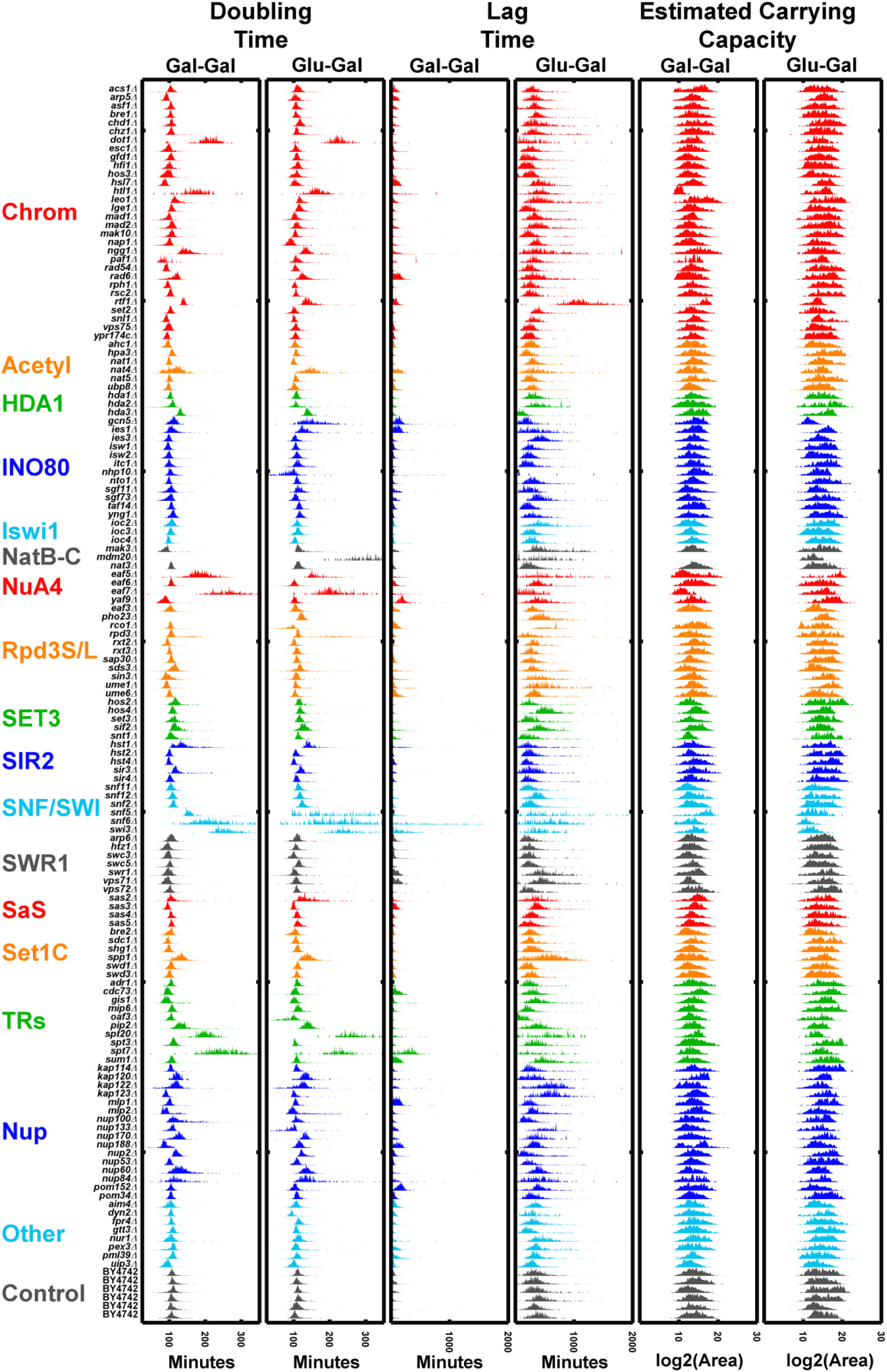
Comparison of 140 Deletion Strains: Doubling times, lag times and estimated carrying capacities of 140 deletion strains that underwent a glucose to galactose switch versus those that were maintained on galactose as a carbon source. Strains are grouped into categories according to annotated gene function (yeastgenome.org) including chromatin modifiers (Chrom), acetyltransferase enzymes (Acetyl), protein complexes (HDA1, INO80, NatB/C,NuA4, Rpd3S/L, Set3, SIR2, SNF/SWI, SWR1, SaS, SetC), transcriptional regulators (TRs), nucleoporins (Nup), and other genes associated with carbon source switching (other).

In general, reporting absolute values of growth parameters is rare in the literature. Here we present a second large-scale application of ODELAY -- to compare doubling times of yeast mutants to the parent strain. A commonly overlooked class of mutant includes the C-terminal tagging with GFP, which is often assumed to have negligible effects on growth when compared with the more dramatic growth defects observed in deletion strains. We tested this assumption by comparing the doubling time of the previously mentioned deletion strains and the corresponding GFP fusion strains against their parent strain, BY4742 (Tables 1 and 2 and supplementary data). All measurements were repeated in triplicate on rich glucose media with the most frequently observed doubling time, the population mode, of each replicate was compared to the parent strain using the Students T-test. ODELAY was able to resolve 12 GFP fusions with doubling times significantly decreased compared to BY4742 and 71 strains that have significantly increased doubling times (Table 1). The deletion strains had 11 strains with significantly decreased doubling times while 72 had significantly increased doubling times (Table 2). While the majority of the doubling time differences for the GFP strains were less than 5 minutes, the presence of the GFP tag does appear to have a wide-spread and significant impact on growth rates on rich media.

**Table 2:** Deletion library strains with significant difference in doubling time over wild type

| Strain |  | Mean<br>Doubling<br>Time<br>Increase | Doubling<br>Time<br>SDEV | P-Val | Strain |  | Mean<br>Doubling<br>Time<br>Increase | Doubling<br>Time<br>SDEV | P-Val |
| --- | --- | --- | --- | --- | --- | --- | --- | --- | --- |
| *snl1Δ* | YIL016W | -7.92 | 0.16 | 8.4E-17 | *sap30Δ* | YMR263W | 2.42 | 1.62 | 3.0E-03 |
| *cdc73Δ* | YLR418C | -3.41 | 1.26 | 3.9E-05 | *gfd1Δ* | YMR255W | 2.49 | 0.40 | 2.0E-03 |
| *asf1Δ* | YJL115W | -3.34 | 2.19 | 7.4E-05 | *nto1Δ* | YPR031W | 2.69 | 0.95 | 9.6E-04 |
| *nat5Δ* | YOR253W | -3.14 | 1.35 | 1.4E-04 | *hos3Δ* | YPL116W | 2.69 | 1.99 | 1.1E-03 |
| *rad54Δ* | YGL163C | -2.65 | 0.99 | 1.1E-03 | *rco1Δ* | YMR075W | 2.71 | 2.09 | 1.0E-03 |
| *swr1Δ* | YDR334W | -2.65 | 0.12 | 1.0E-03 | *nup188Δ* | YML103C | 2.79 | 0.89 | 6.2E-04 |
| *rxt3Δ* | YDL076C | -2.46 | 0.67 | 2.3E-03 | *vps71Δ* | YML041C | 2.87 | 2.14 | 5.6E-04 |
| *dyn2Δ* | YDR424C | -2.04 | 0.80 | 1.1E-02 | *sif2Δ* | YBR103W | 3.04 | 1.50 | 2.2E-04 |
| *acs1Δ* | YAL054C | -2.02 | 1.66 | 1.3E-02 | *loc2Δ* | YLR095C | 3.22 | 1.36 | 1.0E-04 |
| *kap123Δ* | YER110C | -1.72 | 0.39 | 3.0E-02 | *hos4Δ* | YIL112W | 3.69 | 1.73 | 1.1E-05 |
| *sas4Δ* | YDR181C | -1.61 | 0.45 | 4.2E-02 | *snf2Δ* | YOR290C | 4.19 | 2.18 | 1.8E-17 |
| *swc5Δ* | YBR231C | 0.97 | 1.91 | 2.7E-02 | *spp1Δ* | YPL138C | 4.41 | 2.31 | 2.3E-16 |
| *ypr174Δc* | Ypr174c | 1.03 | 1.32 | 3.1E-02 | *lge1Δ* | YPL055C | 4.42 | 1.17 | 2.1E-07 |
| *yng1Δ* | YOR064C | 1.56 | 6.11 | 3.2E-02 | *mak3Δ* | YPR051W | 4.46 | 0.94 | 1.6E-07 |
| *nhp10Δ* | YDL002C | 1.59 | 0.57 | 4.6E-02 | *snf12Δ* | YNR023W | 4.49 | 1.34 | 1.4E-07 |
| *taf14Δ* | YPL129W | 1.59 | 1.24 | 4.7E-02 | *nup60Δ* | YAR002W | 4.53 | 0.58 | 9.7E-08 |
| *pex3Δ* | YDR329C | 1.66 | 1.21 | 3.8E-02 | *nup170Δ* | YBL079W | 4.81 | 0.49 | 2.0E-08 |
| *aim4Δ* | YBR194W | 1.67 | 2.16 | 4.0E-02 | *pho23Δ* | YNL097C | 4.83 | 1.46 | 2.3E-08 |
| *bre2Δ* | YLR015W | 1.68 | 1.48 | 3.7E-02 | *chz1Δ* | YER030W | 5.06 | 1.67 | 7.1E-09 |
| *ubp8Δ* | YMR223W | 1.80 | 0.34 | 2.4E-02 | *itc1Δ* | YGL133W | 5.49 | 0.87 | 3.9E-10 |
| *hos2Δ* | YGL194C | 1.83 | 0.72 | 2.2E-02 | *paf1Δ* | YBR279W | 5.97 | 0.90 | 2.2E-11 |
| *isw1Δ* | YBR245C | 1.83 | 0.66 | 2.1E-02 | *snt1Δ* | YCR033W | 6.59 | 2.49 | 1.5E-12 |
| *sds3Δ* | YIL084C | 1.87 | 0.52 | 1.9E-02 | *asm4Δ* | YDL088C | 6.94 | 8.51 | 1.6E-10 |
| *sum1Δ* | YDR310C | 2.11 | 2.09 | 1.0E-02 | *set3Δ* | YKR029C | 7.55 | 0.52 | 9.7E-16 |
| *sgf73Δ* | YGL066W | 2.21 | 2.44 | 7.5E-03 | *ies3Δ* | YLR052W | 7.57 | 1.53 | 1.5E-15 |
| *swd3Δ* | YBR175W | 2.33 | 3.05 | 5.8E-03 | *swc3Δ* | YAL011W | 7.82 | 2.89 | 1.5E-15 |
| *nup133Δ* | YKR082W | 2.34 | 0.62 | 3.6E-03 | *nup2Δ* | YLR335W | 8.41 | 0.77 | 4.5E-18 |

| Strain |  | Mean<br>Doubling<br>Time<br>Increase | Doubling<br>Time<br>SDEV | P-Val |
| --- | --- | --- | --- | --- |
| *leo1Δ* | YOR123C | 8.45 | 0.85 | 3.7E-18 |
| *gcn5Δ* | YGR252W | 8.47 | 1.63 | 5.9E-18 |
| *sir3Δ* | YLR442C | 9.99 | 0.36 | 2.2E-22 |
| *rpd3Δ* | YNL330C | 10.20 | 1.39 | 1.1E-22 |
| *pip2Δ* | YOR363C | 10.58 | 0.76 | 7.2E-24 |
| *ies1Δ* | YFL013C | 11.08 | 3.03 | 7.5E-24 |
| *hda2Δ* | YDR295C | 11.16 | 2.81 | 3.0E-24 |
| *rtf1Δ* | YGL244W | 11.52 | 3.02 | 6.2E-25 |
| *rad6Δ* | YGL058W | 11.96 | 2.83 | 3.5E-26 |
| *kap120Δ* | YPL125W | 13.76 | 1.97 | 3.9E-31 |
| *nup100Δ* | YKL068W | 16.68 | 2.95 | 1.2E-36 |
| *sas2Δ* | YMR127C | 20.44 | 4.35 | 5.5E-42 |
| *ngg1Δ* | YDR176W | 21.17 | 2.52 | 1.4E-45 |
| *kap122Δ* | YGL016W | 22.07 | 4.32 | 8.3E-45 |
| *hda3Δ* | YPR179C | 22.77 | 1.66 | 5.3E-49 |
| *hst1Δ* | YOL068C | 23.95 | 2.62 | 4.5E-50 |
| *nup84Δ* | YDL116W | 24.82 | 1.41 | 1.8E-52 |
| *htl1Δ* | YCR020W-B | 25.90 | 1.86 | 9.1E-54 |
| *nat4Δ* | YMR069W | 26.32 | 2.95 | 2.5E-53 |
| *eaf5Δ* | YEL018W | 28.32 | 6.34 | 4.0E-66 |
| *spt7Δ* | YBR081C | 36.15 | 5.62 | 1.5E-61 |
| *ume1Δ* | YPL139C | 39.04 | 4.49 | 1.2E-66 |
| *eaf7Δ* | YNL136W | 41.02 | 13.45 | 1.4E-50 |
| *dot1Δ* | YDR440W | 41.67 | 4.65 | 5.4E-69 |
| *swi3Δ* | YJL176C | 54.40 | 4.50 | 4.1E-80 |
| *spt20Δ* | YOL148C | 73.09 | 8.36 | 3.4E-99 |
| *snf6Δ* | YHL025W | 85.92 | 24.82 | 1.4E-60 |
| *snf5Δ* | YBR289W | 101.28 | 70.92 | 1.6E-30 |
| *mdm20Δ* | YOL076W | 101.37 | 10.87 | 1.9E-92 |

ODELAY is a quantitative tool capable of multiparameter growth analysis based on time resolved microcolony expansion on solid media. The unique features of ODELAY include its relatively large dynamic range, when compared to other available methods, which enables quantitative measurement of doubling time, lag time and carrying capacity in a single experiment. Additionally, ODELAY has the ability to assess population heterogeneity, including viability, through the analysis of single microcolonies. These features of ODELAY open unexplored avenues for characterizing cellular fitness for even well studied organisms such as baker’s yeast. Although all initial experiments have utilized haploid baker’s yeast, this methodology can be applied in other colony-forming organisms including medically relevant bacteria such as *Mycobacterium tuberculosis*, *Pseudomonas aeruginosa*, *Staphylococcus aureus* and others.

In this report, we have validated ODELAY by comparison to the well-established OD_600_ liquid culture. These experiments revealed that while both liquid culture OD_600_ assays and ODELAY yield doubling times of similar precision and accuracy, ODELAY has the capability to quantitatively resolve heterogeneity from mixed populations of cells into distinct sub-populations. The examples we depict are mutant members of the SWR1 complex as these genes are involved in chromatin remodeling(Krogan et al., 2003). Heterogeneity in this case is likely indicative of cell dysregulation and therefore expected in the SWR1 complex deletion mutants. However, C-terminal tagging with GFP is often assumed not to induce a growth phenotype yet we clearly observe bimodal growth phenotypes in *ARP6-GFP*. These findings reaffirm that tagging a protein with GFP can influence its function and have consequences that manifest in altered cell regulation and ultimately cell growth. In general, we observe slight but significant reductions in growth rate for many GFP-tagged mutants when grown in rich media and without environmental perturbations. These growth phenotypes may appear with greater magnitude in defined or minimal media. These stratifications of yeast fitness phenotypes promise to add new dimensions for understanding the genetic landscape of the cell. Since small perturbations to genes (such as GFP fusions) are readily observed with ODELAY, the effects of multiple gene perturbations can be statistically quantified and used to investigate genetic regulatory networks unobservable by existing SGA and E-MAP methods.

As with other growth assays, there are caveats associated with ODELAY. Extraction of cell doubling time by ODELAY relies on the assumption that microcolony cross-sectional area is directly proportional to the volume of cells in a given colony and that this relationship between volume and area are unaffected by changes in growth condition and/or genetic background. There will certainly be exceptions to this assumption in yeast, and other colony forming microorganisms; however, similar to the limitations in liquid culture OD_600_ analysis when applied to flocculent mutant strains, such exceptions may yield informative phenotypic information. Furthermore, ODELAY could be adapted to analyze 3D volume of the growing micro-colonies; however, this would trade off time for collecting images or limit the total area interrogated. Lag time measurements were also observed to have local variations, which are also commonly observed in other solid phase growth assays(Baryshnikova et al., 2010; Levin-Reisman et al., 2010). There are several other practical benefits of ODELAY’s high-throughput growth rate analysis on solid media over liquid media assays, including increased sample density and reduced handling and materials. The major benefit of ODELAY is the ease with which to quantitatively monitor, for up to hundreds of thousands of individual microcolonies, growth parameters that together define colony expansion from single seeded cells. ODLEAY provides a resolution beyond that permitted by non-dynamical metrics of fitness.

## ACKNOLEDGEMENTS

The authors acknowledge support for this work by grants U54 RR022220 and P50 GM076547 to J.D.A from the U.S. National Institutes of Health. We also thank the Luxembourg Centre for Systems Biomedicine and the University of Luxembourg for support.

## Author Contributions

D.J.D. conceived the ODELAY method. ODELAY was developed by T.E.H., A.V.R., and D.J.D., with intellectual support from S.L., J.J.S. and J.D.A. T.E.H. and S.L. performed all experiments. T.E.H. and D.J.D. wrote the manuscript with contributions from A.V.R., F.D.M., J.J.S. and J.D.A.

## Supplementary Files

All files are available for download at: http://aitchisonlab.com/ODELAY

1. **ODELAY_README.txt**
2. **ODELAY Strain List.docx The complete list of strains measured in this experiment**
3. **ODELAY MicroscopeControl.zip Files that control Leica DMI6000 microscope with a 10X objective**
4. **ODELAY Image Processing Tool.zip Files that contain a data viewer and data analysis packages**
5. **ODELAY Hardware Design.zip AUTOCAD files and word files that demonstrate how to mold agar media for ODELAY experiments**
6. **ODELAY Stage Mount.zip AUTOCAD files and PDFs that depict the stage assembly utilized to collect ODELAY time course experiments**
7. **ODELAY Sample Data.zip Example data sets for demonstrating ODELAY software functionality.**

---

**Supplementary Table 1.** All strains investigated in this study.

| Strain | Strain | Strain | Strain | Strain | Strain | Strain | Strain |
| --- | --- | --- | --- | --- | --- | --- | --- |
| Common | Systematic | Common | Systematic | Common | Systematic | Common | Systematic |
| Name | Name | Name | Name | Name | Name | Name | Name |
| Acs1 | YAL054C | Hst2 | YPL015C | Nup133 | YKR082W | Shg1 | YBR258C |
| Adr1 | YDR216W | Hst4 | YDR191W | Nup170 | YBL079W | Sif2 | YBR103W |
| Ahc1 | YOR023C | Htl1 | YCR020W-B | Nup188 | YML103C | Sin3 | YOL004W |
| Aim4 | YBR194W | Htz1 | YOL012C | Nup2 | YLR335W | Sir3 | YLR442C |
| Arp5 | YNL059C | Ies1 | YFL013C | Nup53 | YMR153W | Sir4 | YDR227W |
| Arp6 | YLR085C | Ies3 | YLR052W | NUP60 | YAR002W | Snf11 | YDR073W |
| Asf1 | YJL115W | loc2 | YLR095C | NUP84 | YDL116W | Snf12 | YNR023W |
| Bre1 | YDL074C | loc3 | YFR013W | Oaf3 | YKR064W | Snf2 | YOR290C |
| Bre2 | YLR015W | loc4 | YMR044W | Paf1 | YBR279W | Snf5 | YBR289W |
| Cdc73 | YLR418C | Isw1 | YBR245C | Pex3 | YDR329C | Snf6 | YHL025W |
| Chd1 | YER164W | Isw2 | YOR304W | Pho23 | YNL097C | Snl1 | YIL016W |
| Chz1 | YER030W | Itc1 | YGL133W | Pip2 | YOR363C | Snt1 | YCR033W |
| Dot1 | YDR440W | Kap114 | YGL241W | Pml39 | YML107C | Spp1 | YPL138C |
| Dyn2 | YDR424C | Kap120 | YPL125W | Pom152 | YMR129W | Spt20 | YOL148C |
| Eaf3 | YPR023C | Kap122 | YGL016W | Pom34 | YLR018C | Spt3 | YDR392W |
| Eaf5 | YEL018W | Kap123 | YER110C | Rad54 | YGL163C | Spt7 | YBR081C |
| Eaf6 | YJR082C | Leo1 | YOR123C | Rad6 | YGL058W | Sum1 | YDR310C |
| Eaf7 | YNL136W | Lge1 | YPL055C | Rco1 | YMR075W | Swc3 | YAL011W |
| Esc1 | YMR219W | Mad1 | YGL086W | Rpd3 | YNL330C | Swc5 | YBR231C |
| Fpr4 | YLR449W | Mad2 | YJL030W | Rph1 | YER169W | Swd1 | YAR003W |
| Gcn5 | YGR252W | Mak10 | YEL053C | Rsc2 | YLR357W | Swd3 | YBR175W |
| Gfd1 | YMR255W | Mak3 | YPR051W | Rtf1 | YGL244W | Swi3 | YJL176C |
| Gis1 | YDR096W | Mdm20 | YOL076W | Rxt2 | YBR095C | Swr1 | YDR334W |
| Gtt3 | YEL017W | Mip6 | YHR015W | Rxt3 | YDL076C | Taf14 | YPL129W |
| Hda1 | YNL021W | Mlp1 | YKR095W | Sap30 | YMR263W | Ubp8 | YMR223W |
| Hda2 | YDR295C | Mlp2 | YIL149C | Sas2 | YMR127C | Uip3 | YAR027W |
| Hda3 | YPR179C | Nap1 | YKR048C | Sas3 | YBL052C | Ume1 | YPL139C |
| Hfi1 | YPL254W | Nat1 | YDL040C | Sas4 | YDR181C | Ume6 | YDR207C |
| Hos2 | YGL194C | Nat3 | YPR131C | Sas5 | YOR213C | Vps71 | YML041C |
| Hos3 | YPL116W | Nat4 | YMR069W | Sdc1 | YDR469W | Vps72 | YDR485C |
| Hos4 | YIL112W | Nat5 | YOR253W | Sds3 | YIL084C | Vps75 | YNL246W |
| Hpa3 | YEL066W | Ngg1 | YDR176W | Set2 | YJL168C | Yaf9 | YNL107W |
| Hsl7 | YBR133C | Nhp10 | YDL002C | Set3 | YKR029C | Ydl089w | Ydl089w |
| Hst1 | YOL068C | Nto1 | YPR031W | Sgf11 | YPL047W | Yng1 | YOR064C |
|  |  | Nup100 | YKL068W | Sgf73 | YGL066W | Ypr174c | Ypr174c |

